# Comparative Transcriptomics Identifies Potential Stemness-Related Markers for Mesenchymal Stromal/Stem Cells

**DOI:** 10.1101/2021.05.25.445659

**Authors:** Myret Ghabriel, Ahmed El Hosseiny, Ahmed Moustafa, Asma Amleh

## Abstract

Mesenchymal stromal/stem cells (MSCs) are multipotent cells residing in multiple tissues with the capacity for self-renewal and differentiation into various cell types. These properties make them promising candidates for regenerative therapies. MSC identification is critical in yielding pure populations for successful therapeutic applications; however, the criteria for MSC identification proposed by the International Society for Cellular Therapy (ISCT) is inconsistent across different tissue sources. In this study, we aimed to identify potential markers to be used together with the ISCT’s criteria to provide a more accurate means of MSC identification. Thus, we carried out a comparative analysis of the expression of human and mouse MSCs derived from multiple tissues to identify the common differentially expressed genes. We show that six members of the proteasome degradation system are similarly expressed across MSCs derived from bone marrow, adipose tissue, amnion, and umbilical cord. Also, with the help of predictive models, we found that these genes successfully identified MSCs across all the tissue sources tested. Moreover, using genetic interaction networks, we showed a possible link between these genes and antioxidant enzymes in the MSC antioxidant defense system, thereby pointing to their potential role in prolonging the life span of MSCs. Our results suggest that these genes can be used as stemness-related markers.

## Introduction

Mesenchymal stem cells (MSCs) are multipotent adult stem cells that can be isolated from a variety of tissues such as bone marrow (BM)^1^, adipose tissue (AT)^2^, amnion (AM)^3^, umbilical cord (UC)^4^, and many more. Due to the myriad sources of MSCs, the International Society for Cellular Therapy (ISCT) proposed minimum criteria by which these MSCs can be identified. These criteria include 1) plastic-adherence of cells in vitro, 2) expression of specific cell surface markers (CD105, CD90, and CD73) and lack of expression of others (CD45, CD14, CD19, CD34, CD11b, CD79alpha, and HLA-DR), and 3) ability to differentiate into osteoblasts, chondroblasts, and adipocytes in vitro^5^. Unfortunately, growing evidence shows that these criteria are not consistent across different tissues and different species since they define only general functional and morphological characteristics^6^. As a result, scientists have resorted to using additional “stemness” or “stemness-related” genes as markers to aid in the correct identification of MSCs^7^. Proper identification of MSCs is crucial to produce pure populations, which will thereby increase the success of their use in regenerative therapies. MSCs are very attractive tools for regenerative therapies, as their ease of isolation and ability to differentiate into multiple lineages make them ideal candidates for this purpose.

MSCs play a critical role in tissue maintenance, regeneration, and homeostasis *in vivo*^8^. Generally, MSCs remain quiescent, relying on glycolysis to produce energy for their metabolic needs^9^; however, upon tissue injury or loss, MSCs are activated to regenerate the damaged tissue and exit quiescence in favor of a more proliferative state. This highly proliferative state is necessary to maintain the balance between replenishing downstream lineages and replenishing the stem cell pool. As they begin to proliferate, energy demands increase and glycolysis shifts to oxidative phosphorylation; this is accompanied by an increase in the production of reactive oxygen species (ROS)^10^. Oxidative phosphorylation is, indeed, a much more efficient means of generating ATP than glycolysis and can produce up to fifteen times more ATP. However, this is a double-edged sword since excess ROS can impair self-renewal and proliferation of MSCs^11,12^. For this reason, MSCs have a very active antioxidant defense system. It has been demonstrated that MSCs constitutively express high levels of antioxidant enzymes, such as superoxide dismutases, catalases, and glutathione peroxidases^13^. These enzymes repair oxidatively damaged proteins, but some proteins become oxidatively modified or damaged in an irreversible way. The cell has systems in place that recognize and remove these irreversibly damaged proteins and consequently prevent their build-up^14^. One of these systems is the proteasome degradation system, which plays an important role in the degradation of oxidized and damaged proteins, preventing their accumulation and subsequent cellular dysfunction^15^.

The proteasome 26S is a multicatalytic degradation complex composed of a core particle (the 20S) and one or two regulatory particles (19S). The 20S core is made up of four rings, two of which are composed of seven alpha subunits, while the other two rings are composed of seven beta subunits. The 19S regulator is comprised of a base (containing six ATPase and two non-ATPase subunits) and a lid (containing up to 10 non-ATPase subunits)^16^. The proteasome’s main function in the cell is to degrade unneeded or damaged proteins by proteolysis; this can be carried out in either a ubiquitin-dependent manner through the 26S pathway or a ubiquitin-independent manner through the 20S pathway. Recently, the proteasome has gained a lot of attention, and it has been shown to play an essential role in the preservation of the self-renewal and stemness of human MSCs. Kapetanou and colleagues^17^ showed that senescence and loss of stemness in human MSCs are accompanied by a sharp decline in proteasome content and activity. Furthermore, they showed that the expression of some proteasome subunits is possibly affected by pluripotency factors. Taken together, these observations support their hypothesis of a relationship between proteostasis and stem cell function, where proteostasis is critical in maintaining proper protein levels, leading to efficient pluripotency and self-renewal maintenance.

In this study, we carried out a comparative analysis of RNA-Seq data of MSCs derived from different tissues of origin (BM, AT, UC, AM) and different species (human, mouse) to yield a list of common differentially expressed genes (DEGs). We provide evidence that members of the proteasome show similar patterns of expression across all MSC samples. Furthermore, we offer a possible relationship between the proteasome and antioxidant enzymes in protecting MSCs from oxidative stress, hence highlighting their importance in MSC survival. Finally, we demonstrate that six members of the proteasomal degradation system can be used as supplementary stemness-related markers for MSC identification through the use of predictive models.

## Results

### RNA-seq analysis

To visualize the transcriptomic similarities and differences between the samples, we constructed clustering maps using the t-distributed stochastic neighbor embedding (t-SNE) statistical method. The clustering maps showed all human MSC samples clustering together in a distinct cluster apart from tissue-specific cell (TSCs) samples (Fig. 1A). Likewise, the mouse MSC samples clustered together, while TSC samples clustered independently (Fig. 1B). Next, we compared the gene expression of human-derived MSCs against their tissue-specific counterparts and identified 20,973, 24,365, 8,296, and 29,197 DEGs for h_AM_MSCs, h_BM_MSCs, h_AT_MSCs, and h_UC_MSCs, respectively. We constructed a Venn diagram to visualize the common DEGs shared by the MSCs derived from the four different tissue sources. The common DEGs made up 4.6% of the examined DEGs, equivalent to 2,181 common DEGs (Fig 2A). Similarly, for the mouse datasets, we compared the gene expression of mouse-derived MSCs with their tissue-specific counterparts and identified 14,843 DEGs for m_AT_MSCs and 12,783 DEGs for the m_BM_MSCs. Again, we constructed a Venn diagram to visualize the common DEGs shared by the MSCs derived from the two different tissues. We found that the common DEGs made up 78.8% of the examined DEGs, equivalent to 12,178 common DEGs (Fig. 2B). Finally, we compared both lists of common DEGs to determine if the human and mouse MSCs shared any common DEGs. The Venn diagram showed that the two species had 1,583 (13.3%) DEGs in common (Fig. 2C). To further investigate the specific expression patterns of these DEGs, we generated a heatmap for the expression of these 1,583 common DEGs. Interestingly, the heatmap showed three main clusters: a cluster that included all the human MSC samples, a cluster that included all the mouse-derived MSC samples plus m_BM_TSCs and, finally, a cluster that included the rest of the TSC samples. Each of these clusters included subclusters that grouped the triplicate of each tissue type (Fig. 3).

**Figure 1.**
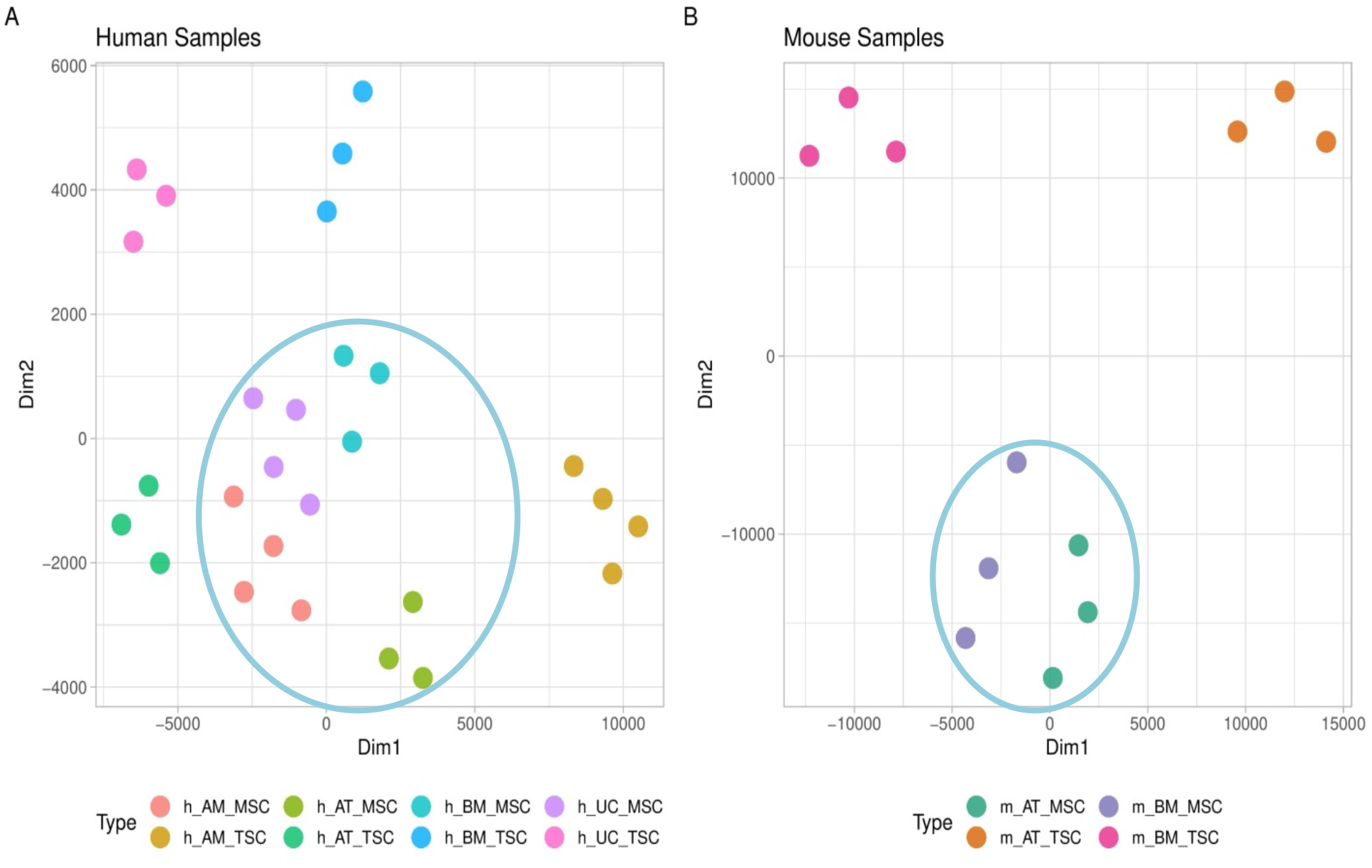
Gene expression-based clustering of MSCs samples included in the study. (A and B) t-SNE clustering shows distinct clusters for human and Mouse MSCs, respectively.

**Figure 2.**
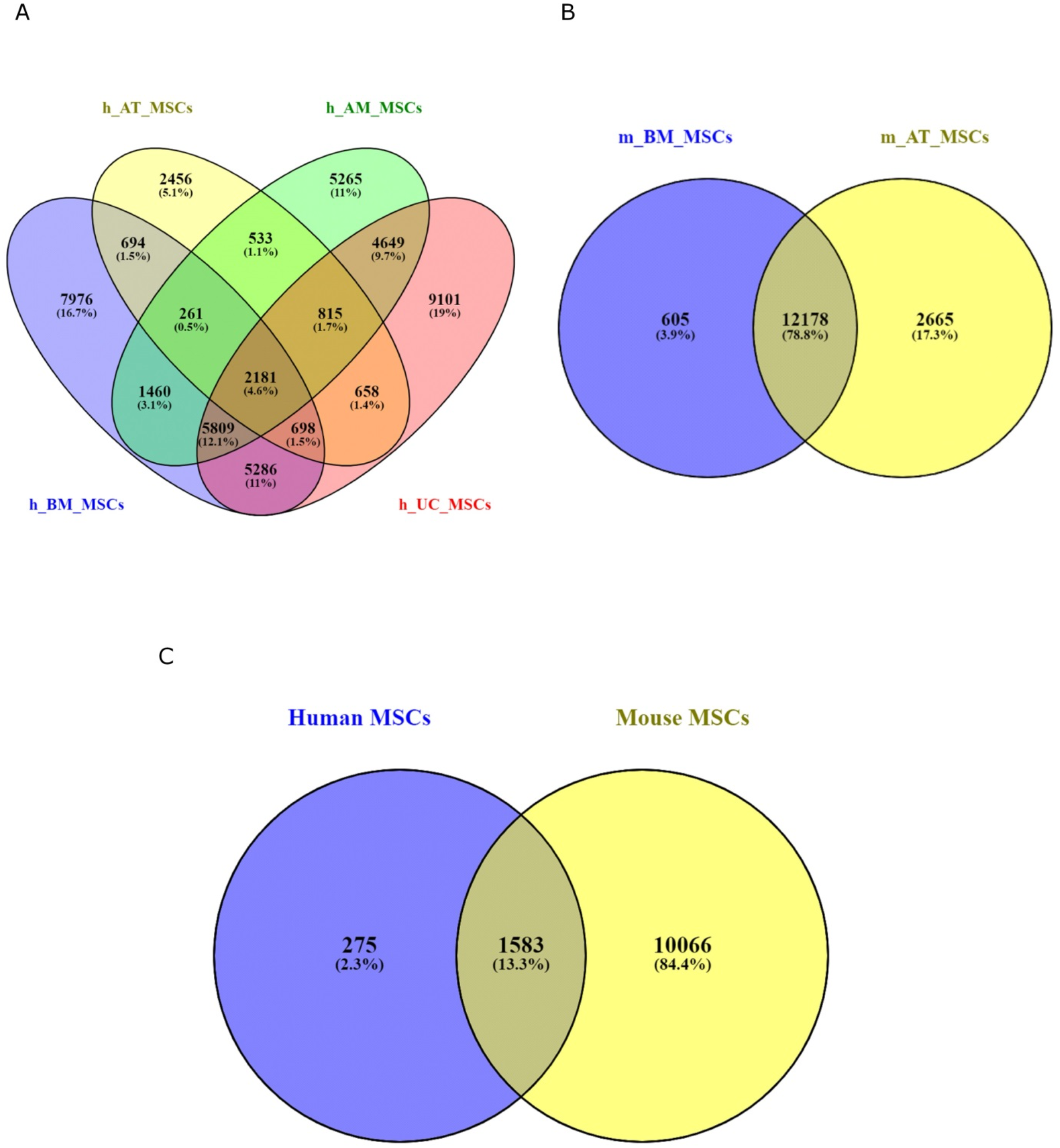
Venn diagram showing the shared and unique DEGs in the transcriptomes of the MSCs derived from different tissues and species. (A) The shared DEGs in the transcriptomes of MSCs derived from four tissue types of human origin are 2181 (4.6%). (B) The shared DEGs between the mouse BM_MSCs and the AT_MSCs are 12178 genes (78.8%). (C) The common DEGs between human and mouse MSCs are1583 (13.3%).

**Figure 3.**
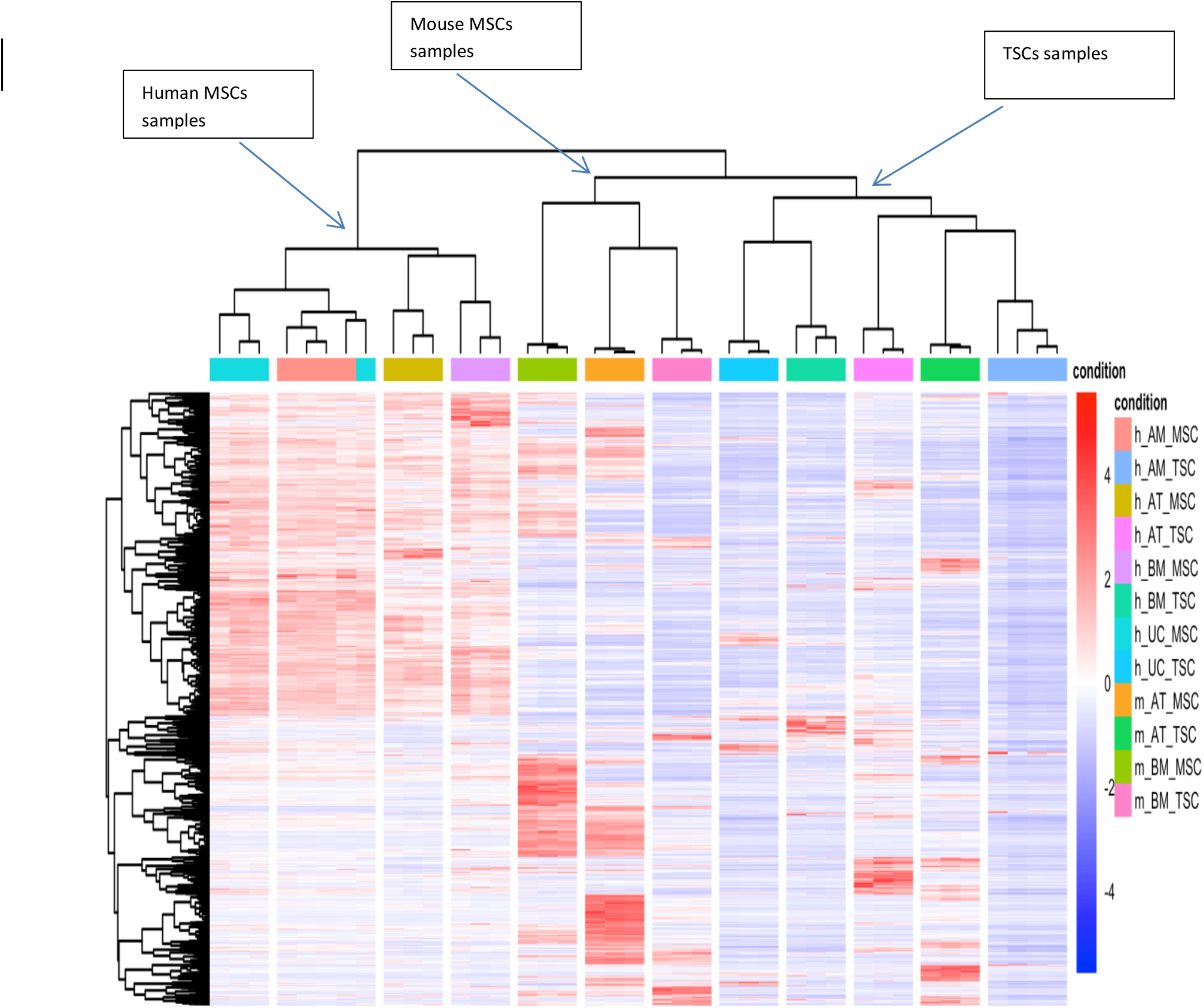
Heatmap of the DEGs across all samples. Heatmap of the 1583 shared MSCs’ DEGs between human and mouse showing three main clusters: a cluster of all human MSCs samples, a cluster of all mouse MSCs samples with m_BM_TSCs, and a cluster of all the TSCs samples.

### DEGs ontology and enrichment analysis

In the gene ontology analysis, we focused on the biological processes–enriched categories to find the processes and pathways most affected by the 1,583 DEGs common to the human and mouse species. We produced a list of 157 enriched gene ontology (GO) terms with a threshold P-value of 10^−3^, which were visualized using ReViGO. ReViGO generated a scatter plot of the enriched GO terms organized according to their significance (p-value) and uniqueness. Following visualization, we identified a unique GO term (GO:2000736) for the regulation of stem cell differentiation with a significant p-value of 8.69E-4, a q value of 4.64E-2, and an enrichment score of 1.48. Moreover, this GO term had a frequency of 0.010%, a log10 p-value of –3.0610, and a high uniqueness score of 0.70 (Fig. 4). Other GO terms were more general, less unique, and not explicitly specific to stem cell function. The GO term GO:2000736 included 23 genes that belonged to the proteasomal degradation pathway (Fig. 5) (Table1).

**Figure 4.**
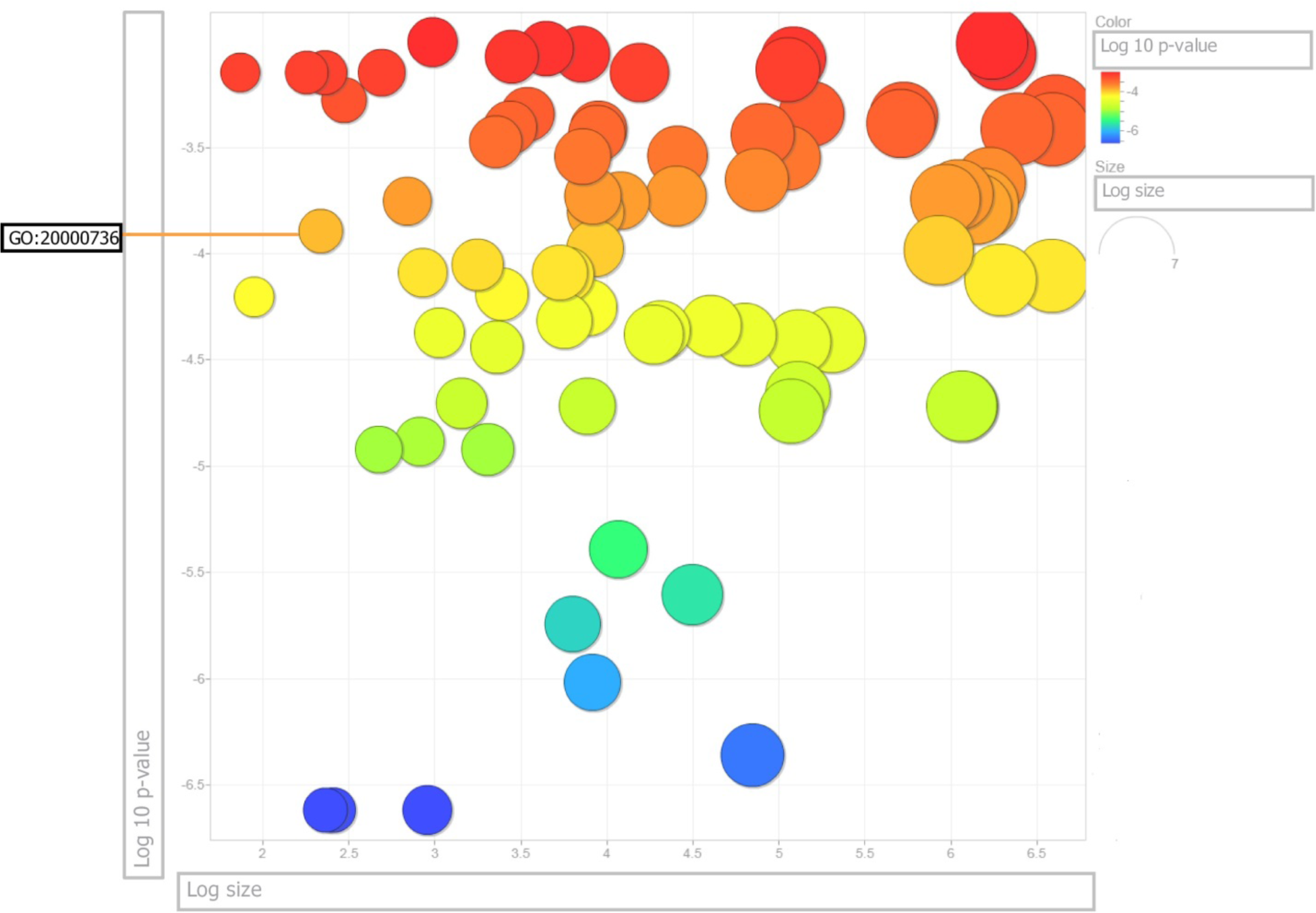
ReViGO scatter plot showing 157 enriched biological processes GO terms associated with the (1583) common DEGs. The color of the circles represents the log10p-value. The orange colour indicates higher significance levels of the GO term enrichment test, while blue indicates low significance. The circle size represents the log size; smaller circles represent more specific GO terms as they are less frequent than general GO terms, which are represented by bigger-sized circles. The GO term GO:2000736 is unique for the regulation of stem cell differentiation and has a frequency of 0.010%, a log10 p-value of −3.0610 and a uniqueness score of 0.70.

**Figure 5.**
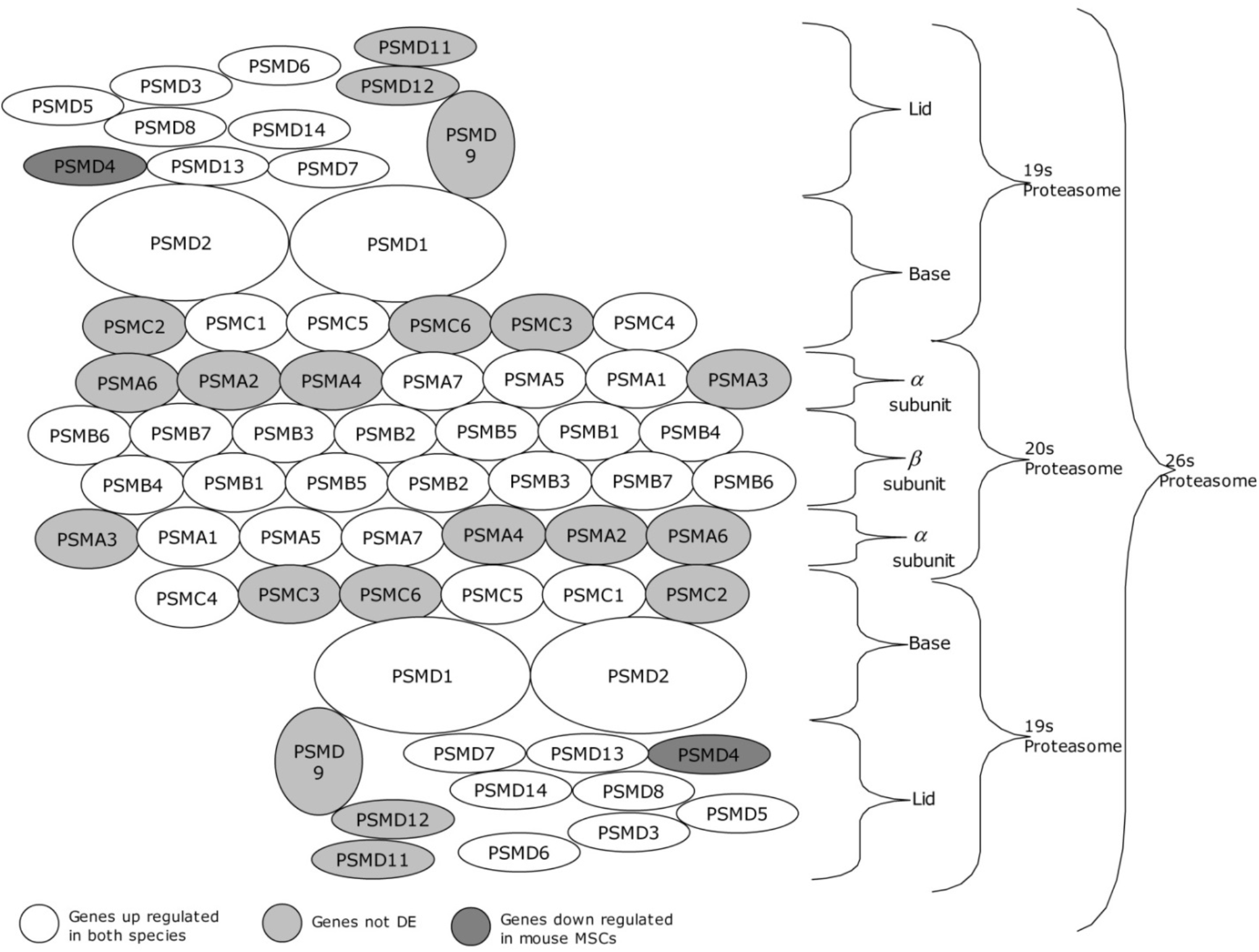
Schematic representation of the different genes involved in the proteasomal degradation machinery shows most are up-regulated. The 26S proteasome is a complex of multiple subunits consisting of 20s and 19s proteasomes. The 20s proteasome is made up of two α-subunits and two β-subunits (each subunit composed of 7 members) while the 19S proteasome is composed of a base and a lid. The light gray colored genes represent the genes not differentially expressed between the MSCs and the TSCs. While the white colored genes represent the twenty two DEGs that are upregulated in MSCs samples across all tissue sources of both human and mouse species. Upregulated DEGs included all seven members of the β –subunit, three members of the α-subunit, five members of the base and seven lid members. The dark gray colored gene represents the only gene downregulated in mouse MSCs.

**Table 1.** List of DEGs belonging to the proteasomal degradation pathway identified in the GO enrichment analysis.

|  | **HUGO**<br>**nomenclature** | **Systemic**<br>**nomenclature** | **Gene name** |
| --- | --- | --- | --- |
| 1 | PSMA1 | $\alpha 6$ | proteasome (prosome, macropain) subunit, alpha type, 1 |
| 2 | PSMA5 | $\alpha 1$ | proteasome (prosome, macropain) subunit, alpha type, 5 |
| 3 | PSMA7 | $\alpha 4$ | proteasome (prosome, macropain) subunit, alpha type, 7 |
| 4 | PSMB1 | $\beta 6$ | proteasome (prosome, macropain) subunit, beta type, 1 |
| 5 | PSMB2 | $\beta 4$ | proteasome (prosome, macropain) subunit, beta type, 2 |
| 6 | PSMB3 | $\beta 3$ | proteasome (prosome, macropain) subunit, beta type, 3 |
| 7 | PSMB4 | $\beta 7$ | proteasome (prosome, macropain) subunit, beta type, 4 |
| 8 | PSMB5 | $\beta 5$ | proteasome (prosome, macropain) subunit, beta type, 5 |
| 9 | PSMB6 | $\beta 1$ | proteasome (prosome, macropain) subunit, beta type, 6 |
| 10 | PSMB7 | $\beta 2$ | proteasome (prosome, macropain) subunit, beta type, 7 |
| 11 | PSMC1 | Rpt2 | proteasome (prosome, macropain) 26s subunit, atpase, 1 |
| 12 | PSMC2 | Rpt1 | proteasome (prosome, macropain) 26s subunit, atpase, 2 |
| 13 | PSMC4 | Rpt3 | proteasome (prosome, macropain) 26s subunit, atpase, 4 |
| 14 | PSMC5 | Rpt6 | proteasome (prosome, macropain) 26s subunit, atpase, 5 |
| 15 | PSMD1 | Rpn2 | proteasome (prosome, macropain) 26s subunit, non-atpase, 1 |
| 16 | PSMD2 | Rpn1 | proteasome (prosome, macropain) 26s subunit, non-atpase, 2 |
| 17 | PSMD3 | Rpn3 | proteasome (prosome, macropain) 26s subunit, non-atpase, 3 |
| 18 | PSMD4 | Rpn10 | proteasome (prosome, macropain) 26s subunit, non-atpase, 4 |
| 19 | PSMD5 | Rpn4 | proteasome (prosome, macropain) 26s subunit, non-atpase, 5 |
| 20 | PSMD7 | Rpn8 | proteasome (prosome, macropain) 26s subunit, non-atpase, 7 |
| 21 | PSMD8 | Rpn12 | proteasome (prosome, macropain) 26s subunit, non-atpase, 8 |
| 22 | PSMD13 | Rpn9 | proteasome (prosome, macropain) 26s subunit, non-atpase, 13 |
| 23 | PSMD14 | Rpn11 | proteasome (prosome, macropain) 26s subunit, non-atpase, 14 |

### Gene expression analysis

Now that our attention was drawn to these 23 genes, we wanted to take a closer look at their behavior. To begin with, we inspected their expression patterns in both species. Heat maps showed that the 23 genes appear to be upregulated in both the human and mouse MSC samples, except for PSMD4, which was downregulated in the mouse MSC samples compared to the TSC samples (Fig. 6A and B). PSMD4 had a P-value of 6.27E-03 and a fold change of 0.136934557 in m_AT_MSCS, while in m_BM_MSCs, it had a P-value of 9.79E-03 and a fold change of –0.224675507. Since most genes were upregulated, we were intrigued by these results and attempted to further explore the interplay between these genes and other systems that assist the proteasome in antioxidant defense, namely, antioxidant enzymes. Consequently, members of the superoxide dismutase, glutathione peroxidase, peroxiredoxin, thioredoxin, and peroxidasin families were cross-referenced against the 1,583 common DEGs identified earlier. We discovered SOD3, GPX7, GPX8, PRDX2, PRDX4, TXN2, and PXDN to be present in our list of common DEGs. We found that the majority of these were upregulated but not all. Out of the seven genes, four (GPX7, GPX8, PXDN, TXN2) were upregulated in all the tested MSC samples. Each of the other three (SOD3, PRDX2, and PRDX4) was found to be upregulated in all the MSC samples except human AT-MSCs and BM-MSCs and mouse BM-MSCs, respectively (Fig. 6C and D).

**Figure 6.**
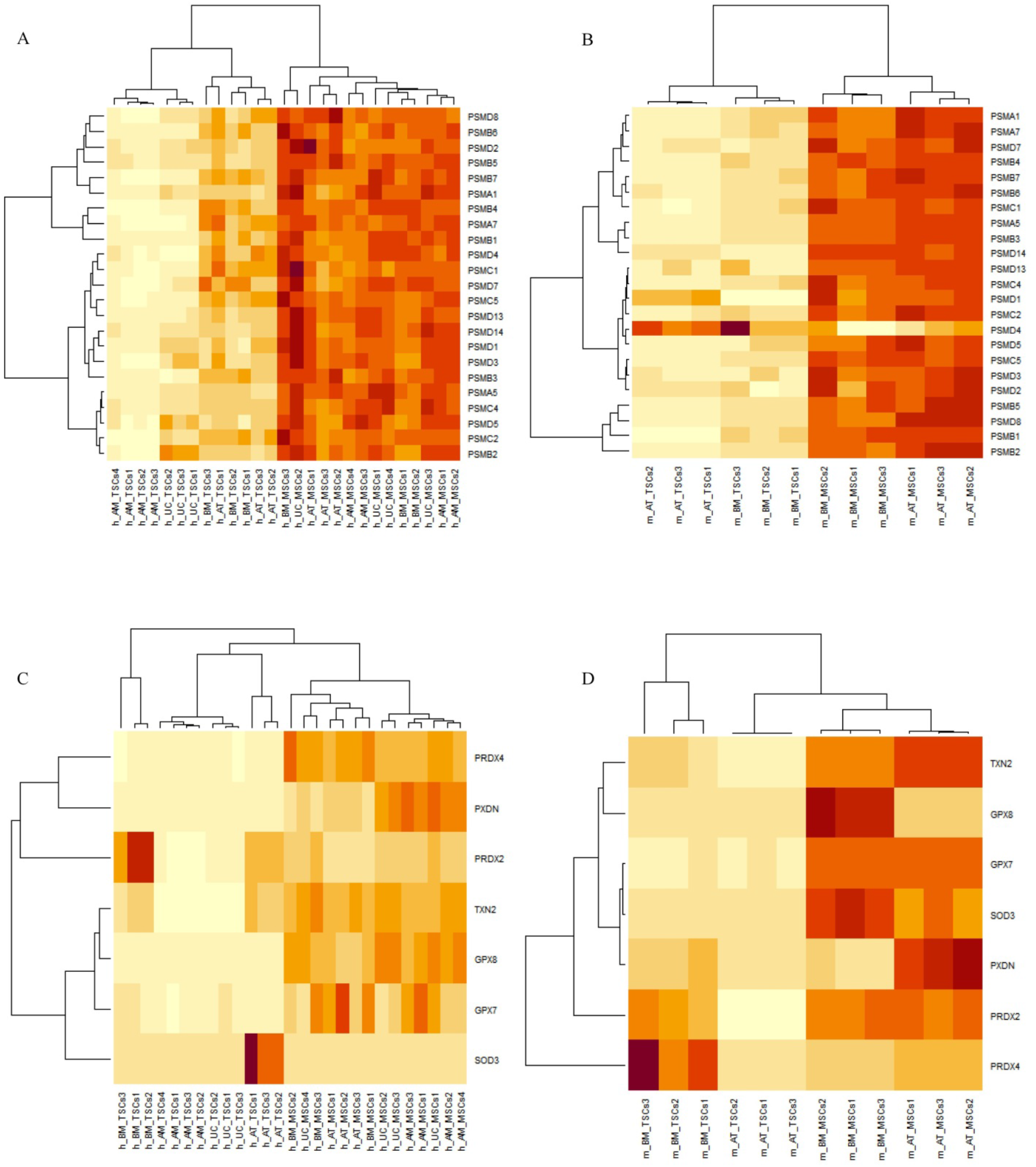
Heatmaps of DEGs expression (rows) in MSCs samples and their tissue-specific counterparts (columns). Orange indicates up-regulation, and yellow indicates down-regulation. (A and B) Heatmaps of the proteasomal genes’ expression in human and mouse samples showing all genes to be upregulated in MSCs samples except PSMD4 in mouse MSCs. (C and D) Heatmaps showing the expression of the antioxidant genes is upregulated in human and mouse MSCs samples with the exception of SOD3 and PRDX2 in human AT_MSCs and BM_MSCs, and PRDX4 in mouse BM_MSCs.

### Gene interaction network

Next, we wanted to shed more light on the interaction between these antioxidant enzymes and the 23 members of the proteasomal degradation system identified earlier. We employed the help of Cytoscape and the Genemania database to understand the interplay between these antioxidant enzymes and the proteasomal genes. A gene interaction network was generated, and it showed all 23 genes of the proteasomal degradation pathway to be co-expressed together and co-expressed with the antioxidant genes. Specifically, it showed PSMA7 to be co-expressed with GPX7, which in turn is co-expressed with GPX8, SOD3, and PXDN. Also, it showed that PRDX2 is co-expressed with PSMA7, PSMB3, PSMB6, PSMB7, PSMC4, PSMD3, and PSMD8. Furthermore, TXN2 was co-expressed with PSMA1, PSMA7, PSMB3, and PSMB6. Finally, PRDX4 was co-expressed with PSMA5, PSMB1, PSMB2, PSMB5, PSMB6, PSMC1, PSMC2, PSMC5, PSMD1, PSMD8, and PSMD14 (Fig. 7).

**Figure 7.**
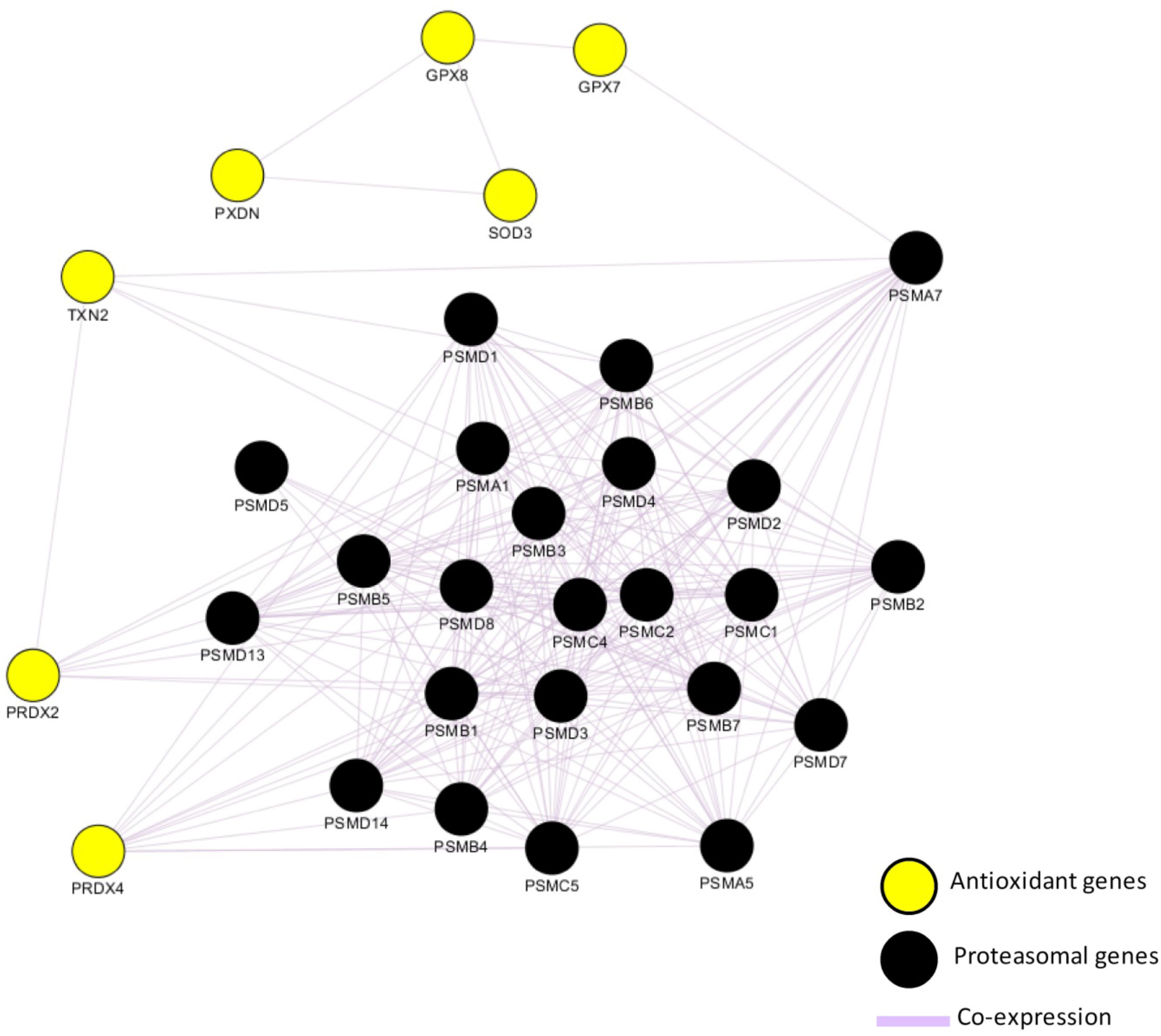
Gene interaction network shows antioxidant genes are co-expressed with proteasomal genes. Cytoscape generated network showing the genes depicted by nodes and the type of interaction depicted by edges. Black circles represent the proteasomal genes that are clearly shown to be co-expressed with the antioxidant genes represented by yellow circles.

### Predictive model

Finally, we wanted to test the genes’ efficiency in predicting the identity of MSCs across the different tissue sources in both human and mouse species. To test this, AutoWEKAClassifier performed 486 evaluations of available classifiers and found random forest to be the best classifier with the best error rate. The random forest tree classifier was used to train the model with 10-fold cross-validation, and the trained model was finalized. The final model was loaded to test its performance in predicting stem cell type on the testing data. We used the upregulated proteasomal genes as attributes, and we removed PSMD4 from the list since it had inconsistent expression across both species. We proceeded with the other 22 genes and ran the random forest model. The model tested the data 40 times and showed that MSCs were correctly classified in all 40 instances. To test if all 22 genes contributed equally to the classification process, we ran the gain ratio attribute selection evaluator in WEKA. We found six genes to be the top contributors in the classification: PSMB5, PSMB1, PSMD14, PSMC4, PSMA1, and PSMD8 (Supplementary Table 3). We repeated the random forest model using these six genes and obtained the same results as the previous model using the 22 genes, and the MSCs correctly classified all 40 instances, as well (Supplementary Table 4).

## Discussion

Ever since the discovery of MSCs by Friedenstein et al.^1^, researchers have debated their identity; however, the criteria proposed by the ISCT still fails to adequately describe MSCs and shows discrepancies across species and tissues of origin. As scientists shift their focus to stemness and stemness-related gene expression to aid in the identification of MSCs, the search for adequate markers has intensified. Here, we show that members of the proteasome degradation system can be used as potential stemness-related markers to identify MSCs.

In this study, we used RNA-seq analysis to aid us in painting a universal portrait for MSC gene expression. To build this comprehensive picture, we integrated the RNA-seq data of MSCs derived from four different tissues (AM, BM, AT, and UC) of human origin and two different tissues (AT, BM) of mouse origin. Differential expression analysis presented us with a list of 1,583 DEGS common to MSCs and TSCs across all tissue types and species. Further gene ontology enrichment analysis categorized these genes into GO terms, one of which was the GO term for regulating stem cell differentiation. This GO term included 23 members of the proteasome degradation system. Compelling evidence suggests a pivotal role for the proteasome in maintaining the pluripotency of mouse and human embryonic stem cells by supporting the clean-up of proteins oxidatively damaged during differentiation^18^.

The proteasome is an essential component of protein quality-control systems and plays an important role in cellular homeostasis. It is involved in the degradation of abnormal, oxidized, or otherwise damaged proteins^19^. The accumulation of oxidized proteins in cells leads to their decreased life span^20^. Moreover, it has been demonstrated that dysfunction of the proteasome is heavily implicated in cell ageing^21^. A recent study revealed that impairment of proteasome function resulted in an accumulation of oxidatively modified proteins in senescent Wharton’s jelly (WJ) and adipose-derived human adult mesenchymal stromal/stem cells. More importantly, this study showed that these cells’ senescence is accompanied by a decline in proteasome content and activities, coupled with the concurrent loss of their stemness^17^. Although the degradation of oxidized proteins can occur by ubiquitin-dependent (26S-proteasome) and ubiquitin-independent (20S-proteasome) mechanisms^22-24^, various studies have shown that the 20S proteasome might be the major machinery involved in this process^25,26^. Here, we show that members of the 20S proteasome (PSMA1, PSMA5, and PSMA7) of the alpha subunits and all members of the beta subunit (PSMB1, PSMB2, PSMB3, PSMB4, PSMB5, PSMB6, and PSMB7) are not only differentially expressed in MSCs but are also upregulated. Furthermore, we show that members of the 19S proteasome base (PSMC1, PSMC2, PSMC4, PSMC5, PSMD1, and PSMD2) and lid (PSMD3, PSMD5, PSMD7, PSMD8, PSMD13, and PSMD14) are also differentially expressed and upregulated in MSCs in comparison with TSCs. However, our results also showed that PSMD4 expression in mouse MSC samples was downregulated. PSMD4’s main role in the 19S lid is to recognize polyubiquitinated protein substrates and to detach the ubiquitin molecules from them for their subsequent degradation through the 26S proteasomal pathway^27^. PSMD4 is not the only ubiquitin receptor in the 19S lid. PSMD2 is another ubiquitin receptor that recognizes and binds both ubiquitin and ubiquitin-like proteins^28^. We found PSMD2 to be upregulated in our mouse MSCs. It could be that mouse MSCs rely mainly on PSMD2 to recognize polyubiquitinated protein substrates, thereby rendering PSMD4 dispensable.

Studies have shown that during MSC proliferation, ROS are produced as byproducts of oxidative metabolism. However, increased ROS levels may lead to a decrease in cell survival and have also been implicated in cell senescence^11^. To counteract these detrimental effects, the cell has antioxidant defense systems that are activated by high ROS levels. Increased ROS concentration causes Nrf2 (a stress-responsive transcription factor) to dissociate from its inhibitory complex with Keap1. This enables Nrf2 to accumulate and translocate to the nucleus, where it binds to antioxidant-response elements (ARE), thus promoting the expression of several antioxidant^29^ and proteasomal genes^30^. These antioxidant enzymes and the proteasome degradation machinery work together as a defense against damaging high ROS levels. Not only did we demonstrate that members of the proteasome degradation machinery were upregulated across the MSCs samples tested, we also demonstrated that GPX7, GPX8, TXN2, and PXDN antioxidant genes were differentially expressed and upregulated in MSCs compared to TSCs. Additionally, by data mining gene interaction databases, we provided evidence that these genes are co-expressed with the proteasome degradation machinery members. Taken together, this points to the efficiency of MSCs in counteracting oxidative stress, in which the proteasome is integral.

Finally, to show the competence of these proteasome genes in identifying MSCs, we employed the aid of predictive models. Predictive models have been used robustly to identify a general MSC phenotype that could distinguish MSCs from other cell types. A recent study showed that gene expression levels in prediction models increase the classification accuracy of the combined set of traditional MSC cell surface markers^31^. Using the random forest model, we showed that the expression of six proteasome genes could accurately distinguish MSCs from their tissue-specific counterparts. Of these six genes, PSMB5, PSMB1, and PSMD14 have been linked to stem cell function. As previously mentioned, PSMB1 and PSMB5 are catalytic subunits of the 20S proteasome, and reducing their expression leads to a decrease in cell proliferation and an increase in replicative senescence in hBMSCs^32^. Likewise, Kapetanou and colleagues reported a similar decrease in the expression of these two genes in senescent WJ-MSCs. They also show that PSMB5 overexpression rescues these senescent cells from age-related decline in proteasome expression and function, thus improving their stemness and extending their lifespan^17^. Finally, PSMD14 is essential for proper 26S assembly^33^; it also plays a role in cleaving polyubiquitin chains at a proximal site and contributes to recycling ubiquitin chains^34^. PSMD14 is a key regulator of stem cell maintenance. A reduction in its levels leads to a marked decrease in Oct4 protein expression, accompanied by abnormal morphology in embryonic stem cells^35^. However, no data currently exists on the three remaining genes (PSMC4, PSMA1, and PSMD8) that link them to any stem cell function.

In our study, we carried out a comprehensive comparative analysis of MSCs RNA-seq data across two species and six different tissue types to ascertain potential identity markers. Our results showed that six members of the proteasomal machinery are promising candidates in identifying MSCs. Moreover, we shine a light on their association with antioxidant enzymes in defending MSCs against high ROS levels, thereby maintaining their proliferation and self-renewal. These six genes can be used as additional stemness-related markers to refine and enhance the accuracy of MSC identification, which is a critical step in ensuring the yield of a pure population for consequent applications in regenerative therapies.

## Materials and methods

### RNA-seq datasets and processing

RNA-seq datasets were obtained from the Gene Expression Omnibus (GEO) ^36^ and Array Express databases^37^. We collected transcriptomic data for human MSCs derived from umbilical cord (h_UC_MSCs), amnion (h_AM_MSCs), bone marrow (h_BM_MSCs), and adipose tissue (h_AT_MSCs) and their tissue-specific counterparts (h_UC_TSCs, h_AM_TSCs, h_BM_TSCs, and h_AT_TSCs). Mouse transcriptomic data included bone marrow-derived MSCs (m_BM_MSCs) and adipose tissue–derived MSCs (m_AT_MSCs), along with their tissue-specific counterparts (m_BM_TSCs and m_AT_TSCs) (Supplementary Table 1). We processed triplicates of each cell type except for h_AM_TSCs, h_AM_MSCs, and h_UC_MSCs, for which we managed to obtain quadruplicates, bringing our total number of samples to 27 human samples and 12 mouse samples (Supplementary Table 2). Data for all cell types were converted from the Sequence Read Archive (SRA) format into the FASTQ format using the SRA Toolkit version 2.10.8 for downstream analysis^38^. Moreover, data was filtered; any read with a length less than 50 bp was excluded. Adapter sequences were detected and trimmed using fastp version 0.19.5^39^.

### RNA-seq data analysis to find DEGs

We used Kallisto version 0.46.1 for pseudo-alignment and the quantification of abundances of transcripts from the RNA-Seq data^40^. Human data was pseudo-aligned to the human reference transcriptome GCA_000001405.15_GRCh38, while mouse data was pseudo-aligned to the mouse reference transcriptome GCA_000001635.9_GRCm39 provided by the Genome Reference Consortium^41^. Abundances produced by Kallisto were processed by Sleuth version 0.29.0 for the differential expression analysis^42^. Venn diagrams of the common DEGs were constructed using Venny version 2.0^43^. Finally, the t-distributed stochastic neighbor-embedding clustering and heatmaps were generated using R 3.6.1 (www.r-project.org).

### DEGs ontology and enrichment analysis

Biological processes encompassing the DEGs were identified based on GO enrichment analysis using the GOrilla database^44^. The *p*-value threshold was set at 10e-3. Afterward, we visualized the enriched GO terms using ReViGO, and a scatter plot was produced showing the log10 p-value and log size of each GO term^45^.

### Generation of gene interaction network

To further investigate the interactions between the DEGs, we constructed a gene interaction network by mining interaction networks from the GEO, BioGRID, IRefIndex, and I2D using the GeneMANIA Cytoscape plugin^46^. This step produced an annotated Cytoscape network of functional interactions between the DEGs.

### Predictive model

Lastly, to assess the selected DEGs’ ability to identify MSCs, we built a predictive model using the Waikato Environment for Knowledge Analysis (WEKA) software version 3.8.4^47^. The gene expression values were converted into the ARFF file format, where our genes of interest were used as attributes in the training dataset. We used the AutoWEKAClassifier package (https://github.com/automl/autoweka) to find the best classification model for our provided dataset automatically. WEKA was also used for attribute selection.

## Author Contribution

Conceived the study: A.A. and M.G. Designed the study: A.A., A.M. and M.G. Collected the data: M.G. and A.E-H. Analyzed the data: M.G., A.E-H., A.M., and A.A. Wrote the paper: M.G., and A.E-H. Review and Editing the paper: A.M., and A.A.

